# HH-suite3 for fast remote homology detection and deep protein annotation

**DOI:** 10.1101/560029

**Authors:** Martin Steinegger, Markus Meier, Milot Mirdita, Harald Vöhringer, Stephan J. Haunsberger, Johannes Söding

## Abstract

**Background:** HH-suite is a widely used open source software suite for sensitive sequence similarity searches and protein fold recognition. It is based on pairwise alignment of profile Hidden Markov models (HMMs), which represent multiple sequence alignments of homologous sequences.

**Results:** We developed a single-instruction multiple-data (SIMD) vectorized implementation of the Viterbi algorithm for profile HMM alignment and introduced various other speed-ups. This accelerated HHsearch by a factor 4 and HHblits by a factor 2 over the previous version 2.0.16. HHblits3 is ~10× faster than PSI-BLAST and ~20× faster than HMMER3. Jobs to perform HHsearch and HHblits searches with many query profile HMMs can be parallelized over cores and over servers in a cluster using OpenMP and message passing interface (MPI). The free, open-source, GNU GPL(v3)-licensed software is available at https://github.com/soedinglab/hh-suite.

**Conclusion:** The added functionalities and increased speed of HHsearch and HHblits should facilitate their use in large-scale protein structure and function prediction, e.g. in metagenomics and genomics projects.

## Introduction

A high sensitivity in sequence similarity searches increases the chance of finding a homologous protein with an annotated function or a known structure from which the function or structure of the query protein can be inferred [1]. Therefore, to find template proteins for comparative protein structure modeling and for deep functional annotation, the most sensitive search tools such as HMMER [2, 3] and HHblits [4] are often used [5–8].

Instead of aligning single sequences with each other, these tools enrich the query sequence using homologous sequences into a multiple sequence alignment (MSA). From the frequencies of amino acids in each column of the MSA, they calculate a 20 × length matrix of position-specific amino acid substitution scores, called sequence profile.

A profile HMM contains, in addition to the position-specific amino acid substitution scores, also position-specific penalties for insertions and deletions relative to the query sequence, which can be estimated from the frequencies of insertions and deletions in the query MSA. The added information improves the sensitivity of profile HMM-based methods like HHblits or HMMER3 over ones based on sequence profiles, such as PSI-BLAST [9].

A few search tools add information on the target side by representing both the query and the target protein by sequence profiles built from MSAs of homologous proteins [10–13]. HHblits / HHsearch represent both the query and the target proteins by profile HMMs. This makes them among the most sensitive tools for sequence similarity search and remote homology detection [4, 14].

In recent years, various sequence search tools have been developed that are up to four orders of magnitude faster than BLAST [15–18]. These tools address the need for faster annotation of massive amounts of environmental sequences being generated through metagenomics against the evergrowing sequence databases. However, no homology can be found for many of these sequences even with sensitive methods, such as BLAST or MMseqs2 [18]. HHblits can help to annotate proteins beyond the twilight zone [19].

In this work, our goal was to accelerate and parallelize various HH-suite algorithms with a focus on the most time-critical tools, HHblits and HHsearch, to facilitate their use on very large datasets. We applied data level parallelization using Advanced Vector Extension 2 (AVX2) or Supplemental Streaming SIMD Extension 3 (SSSE3) instructions, thread level parallelization using OpenMP, and parallelization across computers using MPI. Most important was the ample use of parallelization through SIMD arithmetic units present in all modern Intel, AMD and IBM CPUs, with which we achieved speed-ups per CPU core of a factor 2 to 4.

## Methods

### Overview of HH-suite

The HH-suite software suite contains search tools HHsearch [14] and HHblits [4], HHmake to generate profile HMMs, and various utilities to build databases of MSAs or profile HMMs, to reformat MSAs etc.

HHsearch searches a query profile HMM through a database of target profile HMMs. The search first aligns the query HMM with each of the target HMMs using the Viterbi dynamic programming algorithm, which finds the alignment with the maximum score. The E-value for the target HMM is calculated from the Viterbi score [4]. Target HMMs that reach sufficient significance to be reported are realigned using the Maximum Accuracy algorithm (MAC) [20]. This algorithm maximizes the expected number of correctly aligned pairs of residues minus a penalty between 0 and 1 (parameter `mact`). Values near 0 produce greedy, long, nearly global alignments, values above 0.3 result in shorter, local alignments.

HHblits is an accelerated version of HHsearch that is fast enough to perform iterative searches through millions of profile HMMs, e.g. through the Uniclust profile HMM databases, generated by clustering the UniProt database into clusters of globally alignable sequences [21]. Analogously to PSI-BLAST and HMMER3, such iterative searches can be used to build MSAs by starting from a single query sequence. Sequences from matches to profile HMMs below some E-value threshold (e.g. 10^−3^) are added to the query MSA for the next search iteration.

HHblits has a two-stage prefilter that reduces the number of database HMMs to be aligned with the slow Viterbi HMM-HMM alignment and MAC algorithms. For maximum speed, the target HMMs are represented in the prefilter as discretized sequences over a 219-letter alphabet in which each letter represents one of 219 archetypical profile columns. The two prefilter stages thus perform a profile-to-sequence alignment, first ungapped then gapped, using dynamic programming. Each stage filters away 95% to 99% of target HMMs.

### Overview of changes from HH-suite version 2.0.16 to 3

#### Vectorized Viterbi HMM-HMM alignment

Most of the speed-up was achieved by developing efficient SIMD code and removing branches in the pairwise Viterbi HMM alignment algorithm. The new implementation aligns 4 (using SSSE3) or 8 (using AVX2) target HMMs in parallel to one query HMM.

#### Fast MAC HMM-HMM alignment

We accelerated the Forward-Backward algorithm that computes posterior probabilities for all residue pairs (*i, j*) to be aligned with each other. These probabilities are needed by the MAC alignment algorithm. We improved the speed of the Forward-Backward and MAC algorithms by removing branches at the innermost loops and optimizing the order of indices, which reduced the frequency of cache misses.

#### Memory reduction

We reduced the memory required during Viterbi HMM-HMM alignment by a factor of 1.5 for SSSE3 and implemented AVX2 with only a 1.3 times increase, despite the need to keep scores for 4 (SSSE3) or 8 (AVX2) target profile HMMs in memory instead of just one. This was done by keeping only the current row of the 5 scoring matrices in memory during the dynamic programming (subsection Memory reduction for backtracing and cell-off matrices), and by storing the 5 backtrace matrices, which previously required one byte per matrix cell, in a single backtrace matrix with one byte per cell (subsection From quadratic to linear memory for scoring matrices). We also reduced the memory consumption of the Forward-Backward and MAC alignment algorithms by a factor of two, by moving from storing posterior probabilities with type double to storing their logarithms using type float. In total, we reduced the required memory by roughly a factor 1.75 (when using SSSE3) or 1.16 (when using AVX2).

#### Accelerating sequence filtering and profile computation

For maximum sensitivity, HHmake, HHblits, and HHsearch need to reduce the redundancy within the input MSA by removing sequences that have a sequence identity to another sequence in the MSA larger than a specified cutoff (90% by default) [14]. The redundancy filtering takes time *O*(*NL*^2^), where *N* is the number of MSA sequences and *L* the number of columns. It can be a runtime bottleneck for large MSAs, for example during iterative searches with HHblits. A more detailed explanation is given in subsection SIMD-based MSA redundancy filter.

Additionally, the calculation of the amino acid probabilities in the profile HMM columns from an MSA can become time-limiting. Its run time scales as *O*(*NL*^2^) because for each column it takes a time ~*O*(*NL*) to compute column-specific sequence weights based on the subalignment containing only the sequences that have no gap in that column.

We redesigned these two algorithms to use SIMD instructions and optimized memory access through reordering of nested loops and array indices.

#### Secondary structure scoring

Search sensitivity could be slightly improved for remote homologs by modifying the weighting of the secondary structure alignment score with respect to profile column similarity score. In HH-suite3, the secondary structure score can contribute more than 20% of the total score. This increased the sensitivity to detect remote homologs slightly without negative impact on the high-precision.

#### New features, code refactoring, and bug fixes

HH-suite3 allows users to search a large number of query sequences by parallelizing HHblits/HHsearch searches over queries using OpenMP and MPI (`hhblits_omp` and `hhblits_mpi`, and `hhsearch_omp`). We removed the limit on the maximum number of sequences in the MSAs (option `-maxseqs <max>`). We ported scripts in HH-suite from Perl to Python and added support for the new PDB format mmCIF, which we use to provide precomputed profile HMM and MSA databases for the protein data bank (PDB) [22], Pfam [23], SCOP [24], and clustered UniProt databases (Uniclust) [21].

We adopted a new format for HHblits databases in which the column state sequences used for prefiltering (former *`.cs219` files) are stored in the ffindex format. (The ffindex format was already used in version 2.0.16 for the a3m MSA files and the hhm profile HMM files). This resulted in a ~4s saving for reading the prefilter database and improved scaling of HHblits with the number of cores. We also integrated our discriminative, sequence context-sensitive method to calculate pseudocounts for the profile HMMs, which slightly improves sensitivities for fold-level homologies [25].

To keep HH-suite sustainable and expandable in the longer term, we extensively refactored code by improving code reuse with the help of new classes with inheritance, replacing POSIX threads (pthreads) with OpenMP parallelization, removing global variables, moving from make to cmake, and moving the HH-suite project to GitHub (https://github.com/soedinglab/hh-suite). We fixed various bugs such as memory leaks and segmentation faults occurring with newer compilers.

### Supported platforms and hardware

HHblits is developed under Linux, tested under Linux and macOS, and should run under any Unix-like operating systems. Intel and AMD CPUs that offer AVX2 or at least SSSE3 instruction sets are supported (Intel CPUs: since 2006, AMD: since 2011). PowerPC CPUs with AltiVec vector extensions are also implemented.

Because we were unable to obtain funding for continued support of HH-suite, user support is unfortunately limited to bug fixes for the time being.

### Paralellization by vectorization using SIMD instructions

All modern CPUs possess SIMD units, usually one per core, for performing arithmetic, logical and other operations on several data elements in parallel. In SSE2 and SSSE3, four floating point operations are processed in a single clock cycle in dedicated 128-bit wide registers. Since 2012, the AVX standard allows to process eight floating point operations per clock cycle in parallel, held in 256 bit AVX registers. With the AVX2 extension came support for byte-word- and integer-level operations, e.g. 32 single-byte numbers can be added or multiplied in parallel (32 × 1byte = 256bits). Intel has supported AVX2 since 2013, AMD since 2015.

HHblits 2.0.16 already used SSE2 in its prefilter for gapless and gapped profile-to-sequence alignment processing 16 dynamic programming cells in parallel, but it did not support HMM-HMM alignment using vectorized code.

#### Abstraction layer for SIMD-based vector programming

Intrinsic functions allow to write SIMD parallelized algorithms without using assembly instructions. However, they are tied to one specific variant of SIMD instruction set (such as AVX2), which makes them neither downwards compatible nor future-proof. To be able to compile our algorithms with different SIMD instruction set variants, we implemented an abstraction layer, `simd.h`. In this layer, the intrinsic functions are wrapped by preprocessor macros. Porting our code to a new SIMD standard therefore merely requires us to extend the abstraction layer to that new standard, whereas the algorithm remains unchanged.

The `simd.h` header supports SSSE3, AVX2 and AVX-512 instruction sets. David Miller has graciously extended the `simd.h` abstraction layer to support the AltiVec vector extension of PowerPC CPUs. Algorithm 1 shows a function that computes the scalar product of two vectors.

##### Algorithm 1 Example C code for SIMD abstraction layer

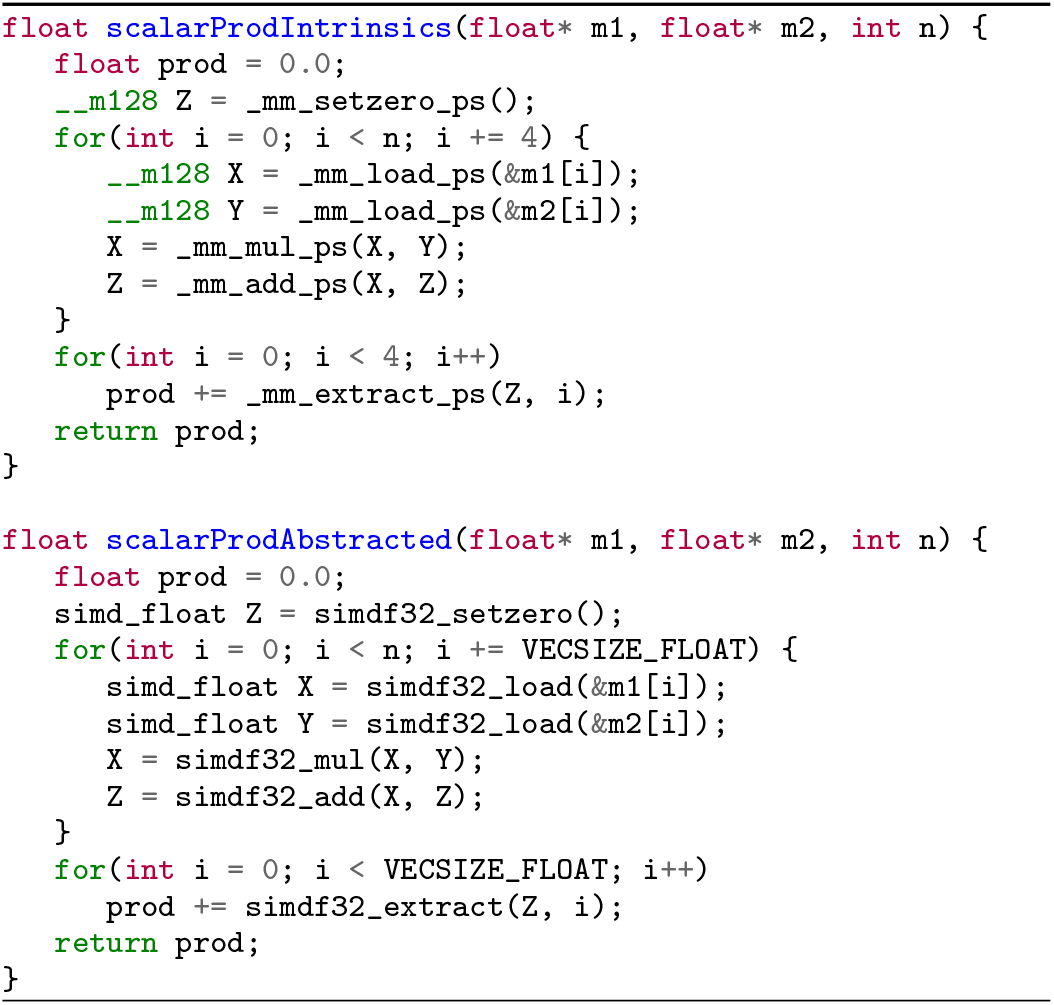

### Vectorized Viterbi HMM-HMM alignments

#### The Viterbi algorithm for aligning profile HMMs

The Viterbi algorithm is a dynamic programming algorithm that generalizes the Smith-Waterman alignment [26]. We use it in HH-suite to compute the best-scoring alignment between two profile HMMs. MSA columns with less than 50% gaps (default value) are modeled by match states in HH-suite, all other columns are modeled as insertion states. By traversing through the states of a profile HMM, the HMM can “emit” sequences. A match state (M) emits amino acids according to the 20 probabilities of amino acids estimated from their fraction in the MSA column, plus some pseudocounts. Insert states (I) emit amino acids according to a standard amino acid background distribution, while delete states (D) do not emit any amino acids.

The alignment score between two HMMs in HH-suite is the sum over all co-emitted sequences of the log odds scores for the probability for the two aligned HMMs to co-emit this sequence divided by the probability of the sequence under the background model. Since M and I states emit amino acids and D states do not, M and I in one HMM can only be aligned with M or I states in the other HMM. Conversely, a D state can only be aligned with a D state or with a Gap G (1). The co-emission score can be written as the sum of the similarity scores of the aligned profile columns, in other words the match-match (MM) pair states, minus the position-specific penalties for indels: delete-open, delete-extend, insert-open and insert-extend.

We denote the alignment pair states as MM, MI, IM, II, DD, DG, and GD. Figure 1 shows an example of two aligned profile HMMs. In the third column HMM *q* emits a residue from its M state and HMM *p* emits a residue from the I state. The pair state for this alignment column is MI. In column six of the alignment HMM *q* does not emit anything since it passes through the D state. HMM *p* does not emit anything either since it has a gap in the alignment. The corresponding pair state is DG. For simplicity and better speed, we exclude pair states II and DD, and we only allow transitions between a pair state and itself and between pair state MM and pair states MI, IM, DG, or GD.

**Figure 1.**
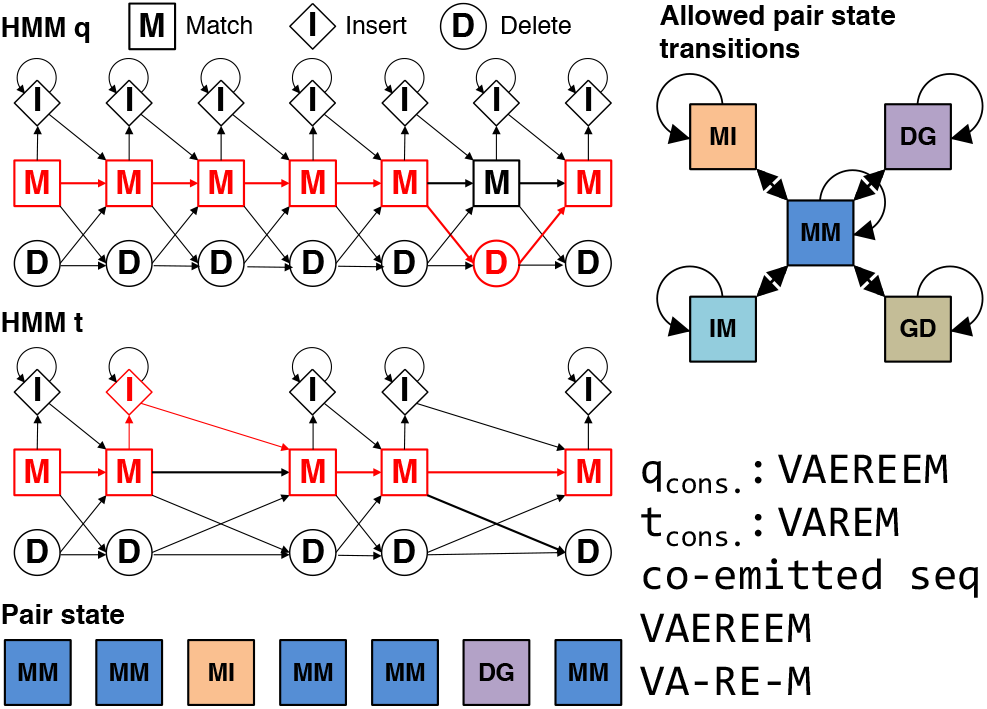
HMM-HMM alignment of query and target. The alignment is represented as red path through both HMMs. The corresponding pair state sequence is MM, MM, Ml, MM, MM, DG, MM.

To calculate the alignment score, we need five dynamic programming matrices *S_XY_*, one for each pair state XY ∈ {MM, MI, IM, DG, GD}. They contain the score of the best partial alignment which ends in column *i* of *q* and column *j* of *p* in pair state XY. These five matrices are calculated recursively.

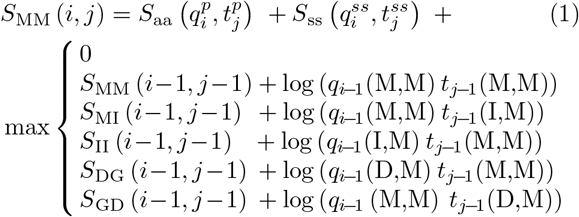

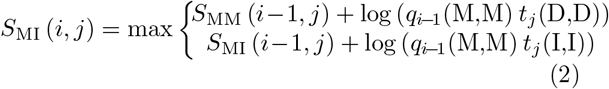

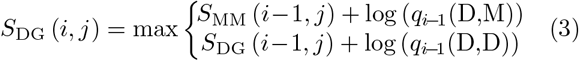

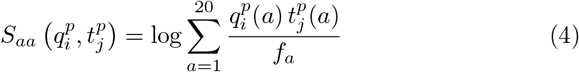

Vector 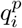 contains the 20 amino acid probabilities of *q* at position *i*, 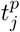 are the amino acid probabilities *t* at *j*, and *f_a_* denotes the background frequency of amino acid *a*. The score *S_aa_* measures the similarity of amino acid distributions in the two columns *i* and *j. S_ss_* can optionally be added to *S_aa_*. It measures the similarity of the secondary structure states of query and target HMM at *i* and *j* [14].

#### Vectorizations of Smith-Waterman sequence alignment

Much effort has gone into improving the performance of the dynamic programming based Smith-Waterman algorithm. While substantial accelerations using general purpose graphics processing units (GPGPUs) and field programmable gated arrays (FPGAs) were demonstrated [27–30], the need for a powerful GPGPU and the lack of of a single standard (e.g. Nvidia’s proprietary CUDA versus the OpenCL standard) have been impediments. SIMD implementations using the SSSE3 and AVX2 standards with on-CPU SIMD vector units have demonstrated similar speed-ups as GPGPIJ implementations and have become widely used [2, 3, 31–34].

To speed up the dynamic programming (DP), the values of several cells in the DP matrix are held in a single SIMD) register and are computed jointly in parallel. Because the value in cell (*i, j*) depends on those in cells (*i* − 1, *j* − 1), (*i* − 1, *j*), and (*i, j* − 1), the cells to be computed at the same time must only depend on cells that have already been computed, not on the cells to be computed at the same time.

Four main approaches have been developed to address this challenge: (1) parallelizing over anti-diagonal stretches of cells in the DP matrices ((*i, j*), (*i* + 1, *j* − 1), … (*i* + 15, *j* − 15), assuming 16 cells fit into one SIMD register) [31], (2) parallelizing over vertical or horizontal segments of the DP matrices (e.g. (*i, j*), (*i* + 1, *j*),… (*i* + 15, *j*)) [32], (3) parallelizing over stripes of the DP matrices ((*i, j*), (*i* + 1 × *D, j*),… (*i* + 15 × *D, j*) where *D* := `ceil`(query_length/16)) [33] and (4) where 16 cells (*i, j*) of 16 target sequences are processed in parallel [34].

##### Algorithm 2 Viterbi algorithm for HMM-HMM alignment

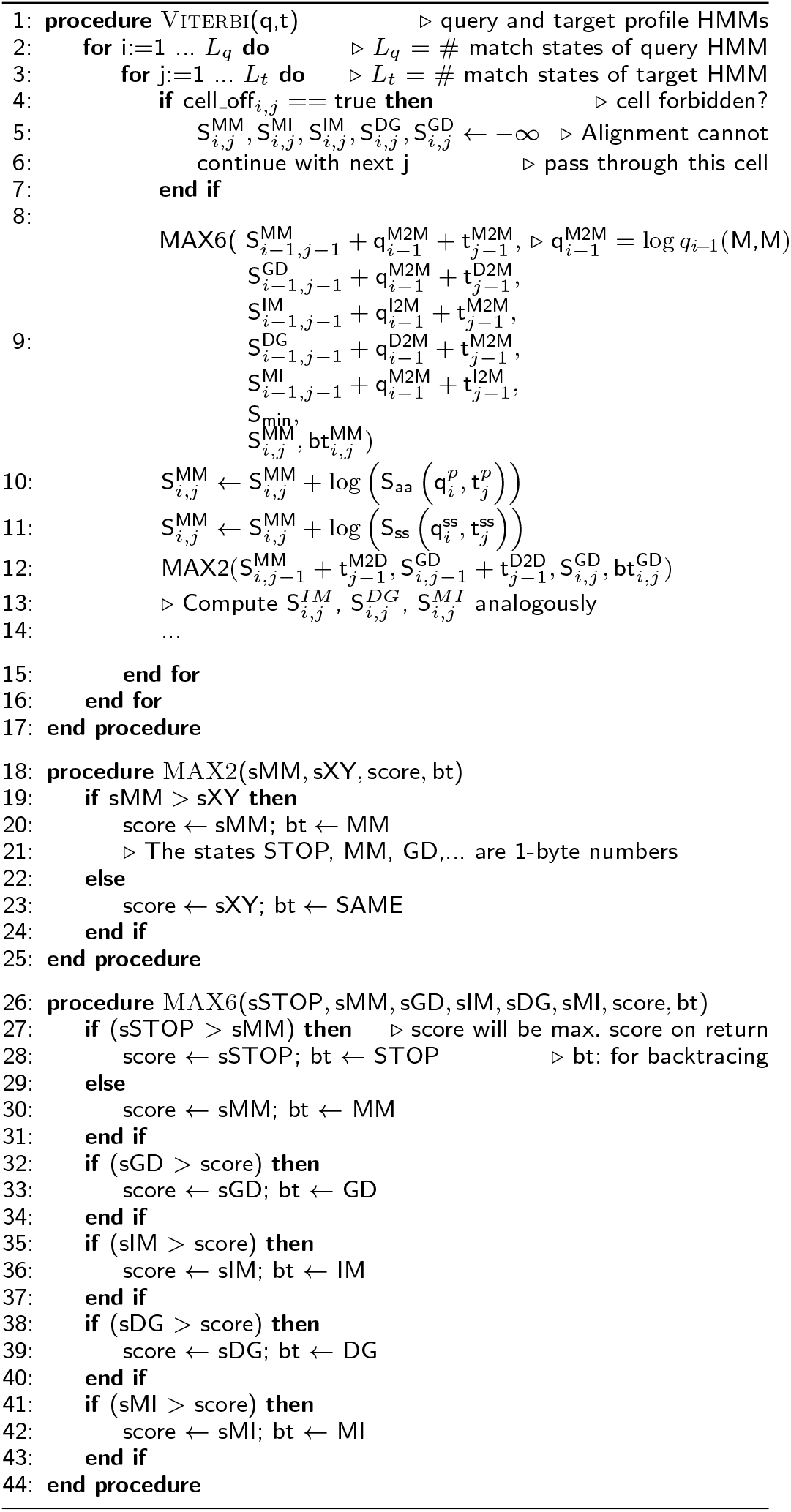

##### Algorithm 3 Branchless, vectorized Viterbi algorithm for HMM-HMM alignment

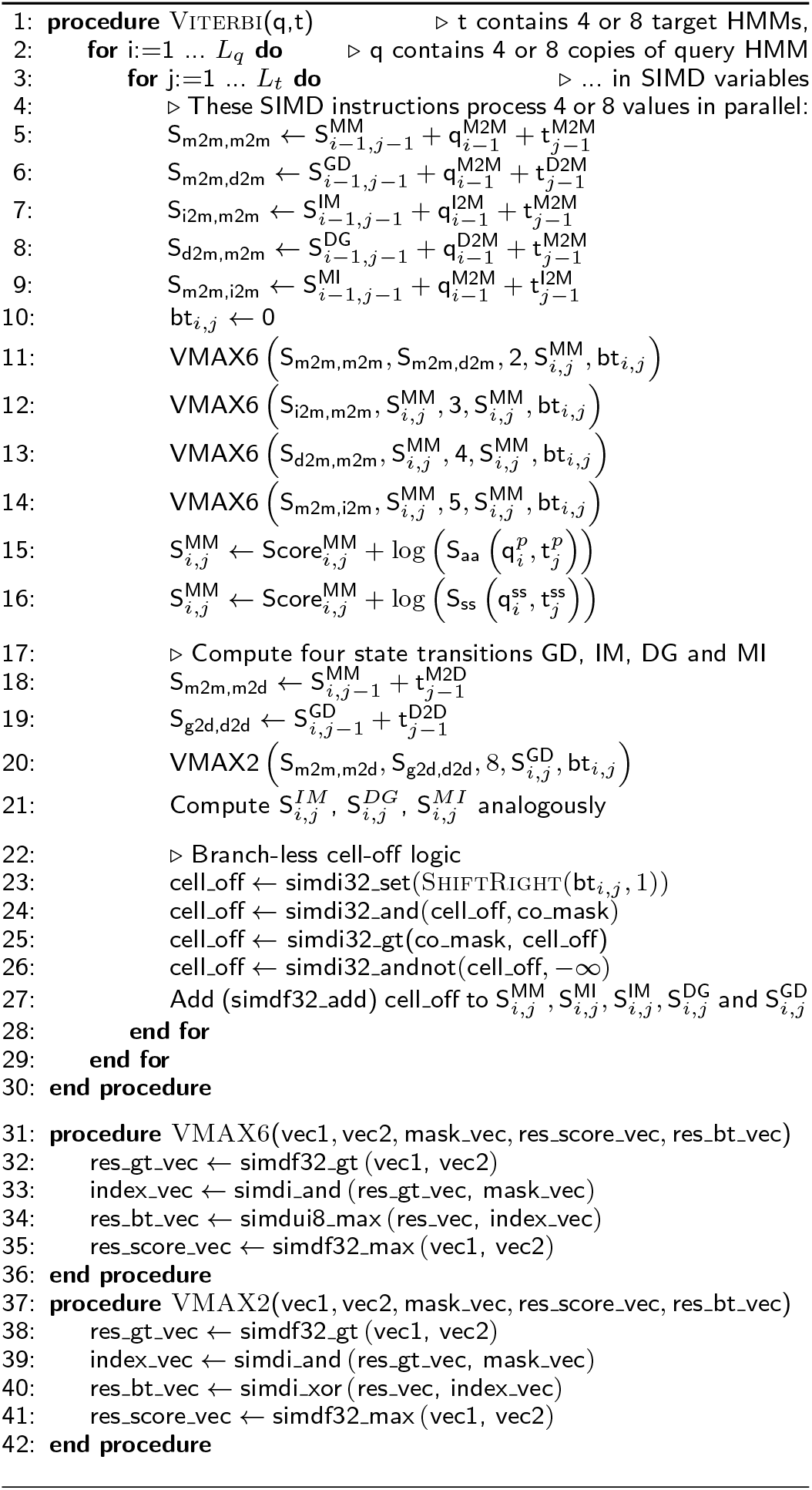

The last option is the fastest method for sequence-sequence alignments, because it avoids data dependencies. Here we present an implementation of this option that can align one query profile HMM to 4 (SSSE3) or 8 (AVX2) target profile HMMs in parallel.

#### Vectorized Viterbi algorithm for aligning profile HMMs

Algorithm 2 shows the scalar version of the Viterbi algorithm for pairwise profile HMM alignment based on the iterative update equations (1)–(3). Algorithm 3 presents our vectorized and branch-less version (Fig. 2). It aligns batches of 4 or 8 target HMMs together, depending on how many scores of type `float` fit into one SIMD register (4 for SSSE3, 8 for AVX).

**Figure 2.**
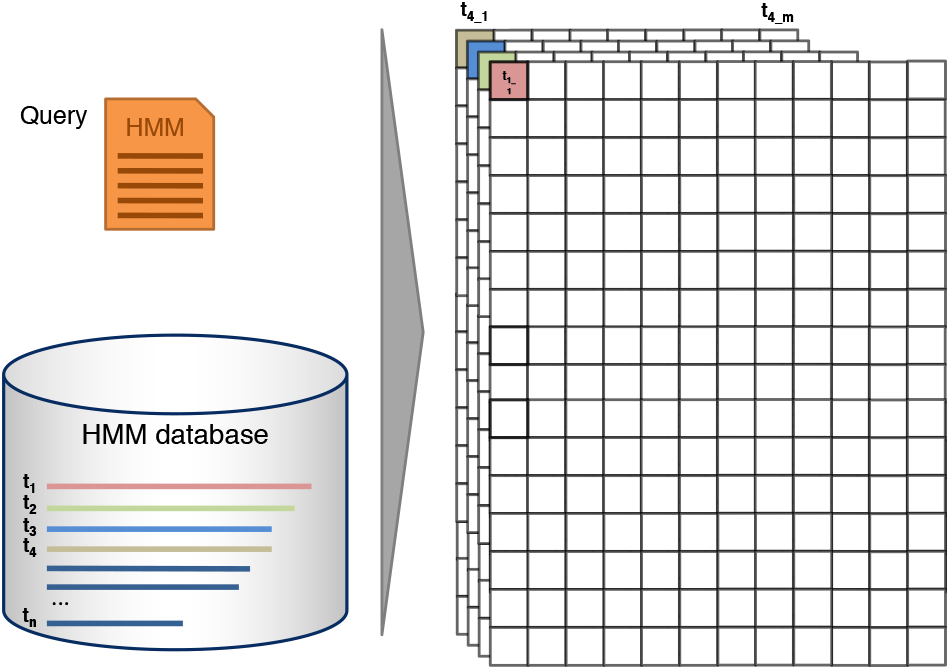
SIMD parallelization over target profile HMMs. Batches of 4 or 8 database profile HMMs are aligned together by the vectorized Viterbi algorithm. Each cell (*i, j*) in the dynamic programming matrix is processed in parallel for 4 or 8 target HMMs.

The vectorized algorithm needs to access the state transition and amino acid emission probabilities for these 4 or 8 targets at the same time. The memory is laid out (Figure 3), such that the emission and transition probabilities of 4 or 8 targets are stored consecutively in memory. In this way, one set of 4 or 8 transition probabilities (for example MM) of the 4 or 8 target HMMs being aligned can be loaded jointly into one SIMD register.

**Figure 3.**
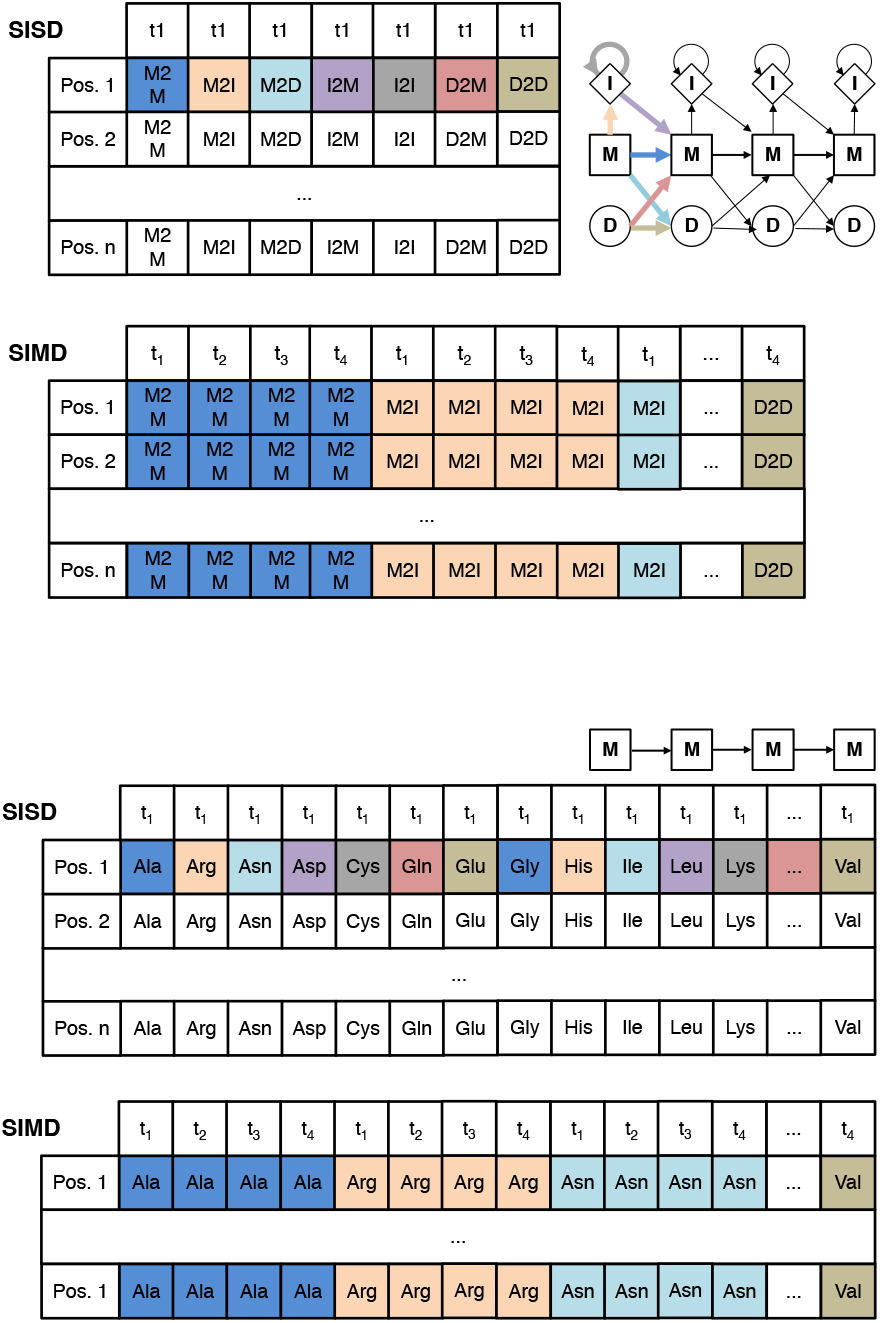
The layout of the log transition probabilities (top) and emission probabilities (bottom) in memory for single-instruction single data (SISD) and SIMD algorithms. For the SIMD algorithm, 4 (using SSSE3) or 8 (using AVX) target profile HMMs (tl − t4) are stored together in interleaved fashion: the 4 or 8 transition or emission values at position *i* in these HMMs are stored consecutively (indicated by the same color). In this way, a single cache line read of 64 bytes can fill four SSSE3 or two AVX2 SIMD registers with 4 or 8 values each.

The scalar versions of the functions `MAX6, MAX2` contain branches. Branched code can considerably slow down code execution due to the high cost of branch mispredictions, when the partially executed instruction pipeline has to be discarded to resume execution of the correct branch.

The functions `MAX6` and `MAX2` find the maximum score out of two or six input scores and also return the pair transition state that contributed the highest score. This state is stored in the backtrace matrix, which is needed to reconstruct the best-scoring alignment once all five DP matrices have been computed.

To remove the five if-statement branches in `MAX6`, we implemented a macro `VMAX6` that implements one if-statement at a time. `VMAX6` needs to be called 5 times, instead of just once as `MAX6`, and each call compares the current best score with the next of the 6 scores and updates the state of the best score so far by maximization. At each `VMAX6` call, the current best state is overwritten by the new state if it has a better score.

We call the function `VMAX2` four times to update the four states GD, IM, DG and MI. The first line in `VMAX2` compares the 4 or 8 values in SIMD register sMM with the corresponding values in register sXY and sets all bits of the four values in SIMD register `res_gt_vec` to 1 if the value in sMM is greater than the one in sXY and to 0 otherwise. The second line computes a bit-wise AND between the four values in `res_gt_vec` (either 0×00000000 or 0×FFFFFFFF) and the value for state MM. For those of the 4 or 8 sMM values that were greater than the corresponding sXY value, we obtain state MM in `index_vec`, for the others we get zero, which represents staying in the same state. The backtrace vector can then be combined using an XOR instruction.

In order to calculate suboptimal, alternative alignments, we forbid the suboptimal alignment to pass through any cell *(i,j)* that is nearer than 40 cells from any of the cells of the better-scoring alignments. These forbidden cells are stored in a matrix `cell_off[i][j]` in the scalar version of the Viterbi algorithm. The first if-statement in Algorithm 2 ensures that these cells obtain a score of −∞.

To reduce memory requirements in the vectorized version, the cell-off flag is stored in the most significant bit of the backtracing matrix (Fig. 5) (see section Memory reduction for backtracing and cell-off matrices). In the SIMD Viterbi algorithm, we shift the backtracing matrix cell-off bit to the right by one and load four `32bit` (SSSE3) or eight 64bit (AVX2) values into a SIMD register (line 23). We extract only the cell-off bits (line 24) by computing an AND between the `co_mask` and the `cell_off` register. We set elements in the register with cell_off bit to 0 and without to `0xFFFFFFFF` by comparing if `cell_mask` is greater than `cell_off` (line 25). On line 26, we set the 4 or 8 values in the SIMD register `cell_off` to −∞ if their cell-off bit was set and otherwise to 0. After this we add the generated vector to all five scores (MM, MI, IM, DG and GD).

A small improvement in runtime was achieved by compiling both versions of the Viterbi method, one with and one without cell-off logic. For the first, optimal alignment, we call the version compiled without the cell off logic and for the alternative alignments the version with cell-off logic enabled. In C/C++, this can be done with preprocessor macros.

Shorter profile HMMs are padded with probabilities of zero up to the length of the longest profile HMM in the batch (Fig. 2). Therefore, the database needs to be sorted by decreasing profile HMM length. Sorting also improves IO performance due to linear access to the target HMMs for the Viterbi alignment, since the list of target HMMs that passed the prefilter is automatically sorted by length.

#### Vectorized column similarity score

The sum in the profile column similarity score *S_aa_* in the first line in Algorithm 4 is is computed as the scalar product between the precomputed 20-dimensional vector 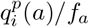 and 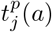. The SIMD code takes 39 instructions to compute the scores for 4 or 8 target columns, whereas the scalar version needed 39 instructions for a single target column.

### From quadratic to linear memory for scoring matrices

Most of the memory in Algorithm 2 is needed for the five score matrices for pair states MM, MI, IM, GD and DG. For a protein of 15 000 residues, the five matrices need 15 000 × 15 000 × 4 byte × 5 matrices = 4.5 GB of memory per thread.

In a naive implementation, the vectorized algorithm would need a factor of 4 or 8 more memory than that, since it would need to store the scores of 4 or 8 target profile HMMs in the score matrices. This would require 36 GB of memory per thread, or 576 GB for commonly used 16 core servers.

However, we do not require the entire scoring matrices to reside in memory. We only need the backtracing matrices, the position (*i*_best_, *j*_best_) and score of the highest scoring cell seen so far to to reconstruct the alignments.

We implemented two approaches. The first uses two vectors per pair state (Fig. 4 top). One holds the scores of the current row *i*, where (*i, j*) are the positions of the cell whose scores are to be computed, and the other vector holds the scores of the previous row *i* − 1. After all the scores of a row *i* have been calculated, the pointers to the vectors are swapped and the former row becomes the current one.

#### Algorithm 4 The similarity scores (eq. (4)) for 4 or 8 target HMMs can be computed in parallel by 39 SIMD vector instructions in just 39 CPU clock cycles

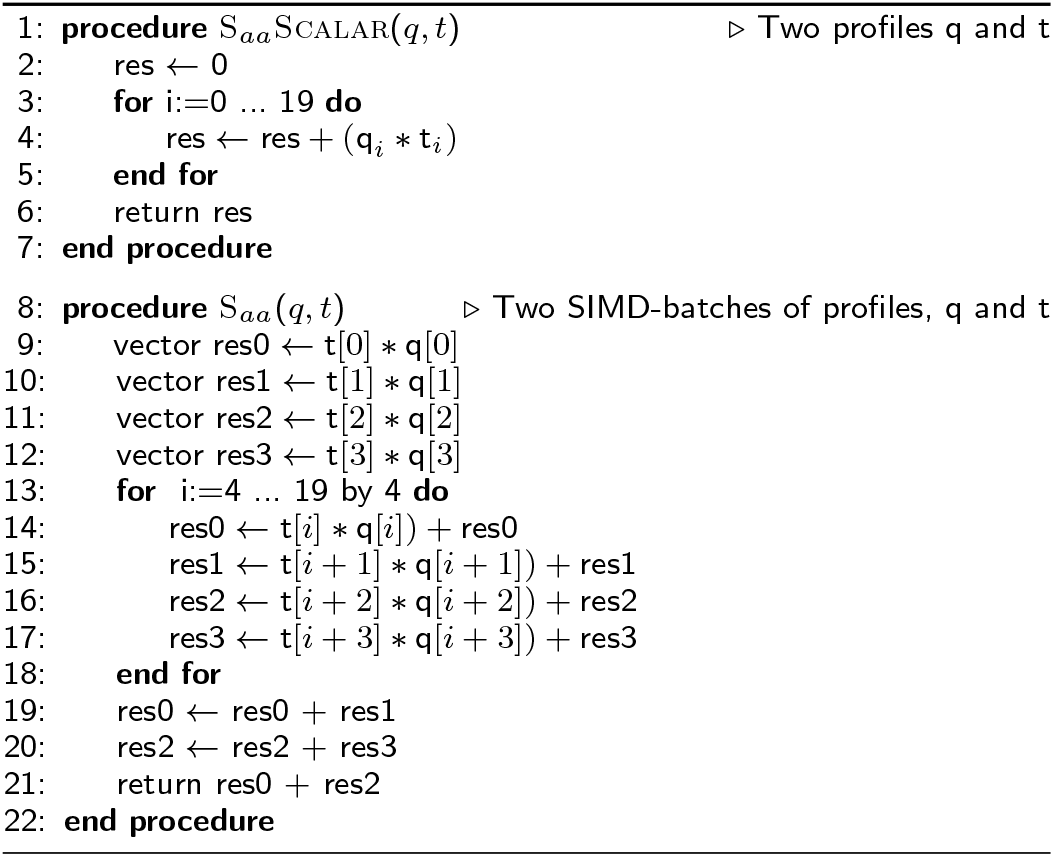

**Figure 4.**
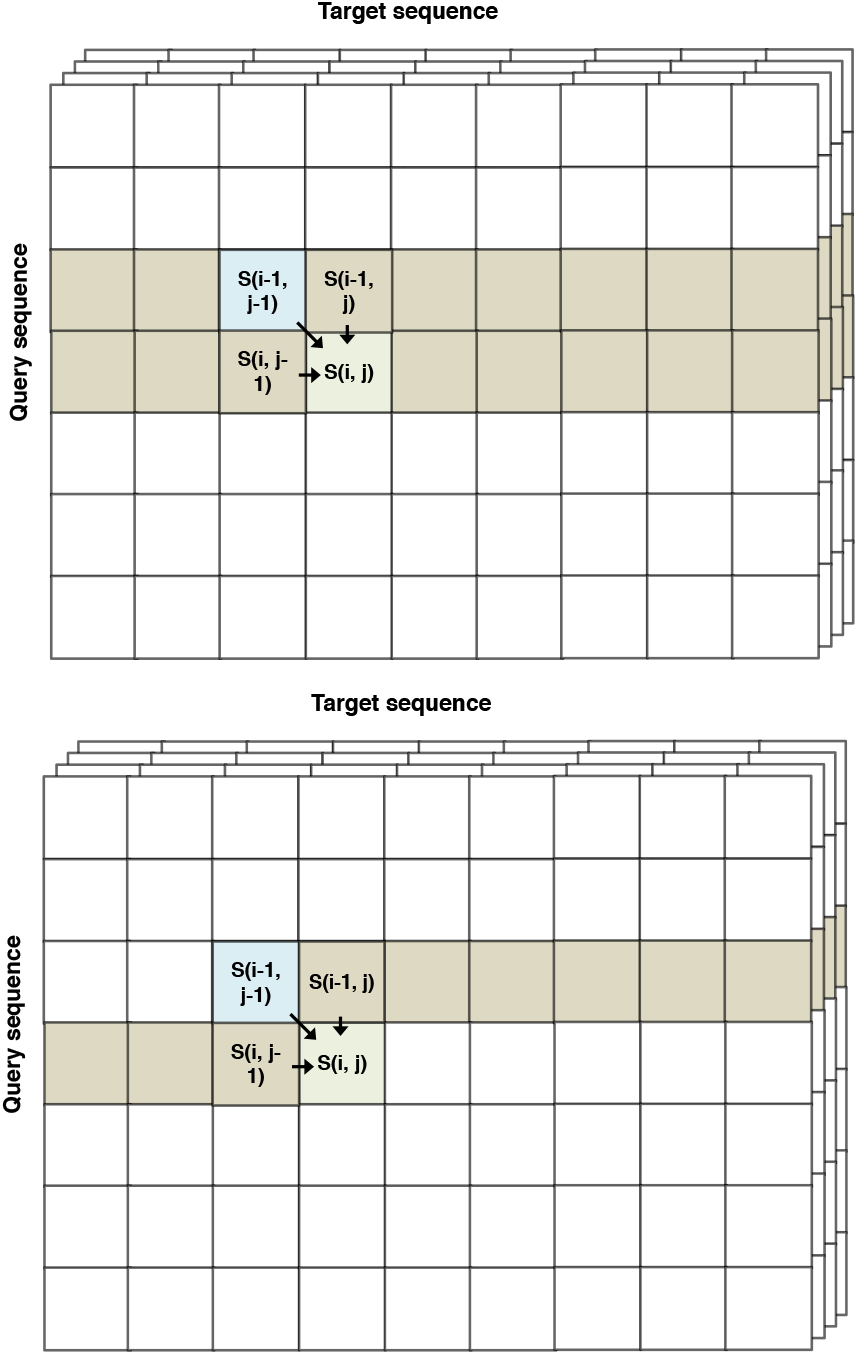
Two approaches to reduce the memory requirement for the DP score matrices from *O*(*L_q_L_t_*) to *O*(*L_t_*), where *L_q_* and *L_t_* are lengths of the query and target profile, respectively. (Top) One vector holds the scores of the previous row, *S*_XY_(*i* − 1, ·), for pair state XY ∈{MM, Ml, IM, GD and DG}, and the other holds the scores of the current row, *S*_XY_(*i* − 1, ·) for pair state XY ∈{MM, Ml, IM, GD and DG}. Vector pointers are swapped after each row has been processed. (Bottom) A single vector per pair state XY holds the scores of the current row up to *j* − 1 and of the previous row for *j* to *L_t_*. The second approach is somewhat faster and was chosen for HH-suite3.

The second approach uses only a single vector (Fig. 4 bottom). Its elements from 1 to *j* − 1 hold the scores of the current row that have already been computed. Its elements from *j* to the last position *L_t_* hold the scores from the previous row *i* − 1.

The second variant turned out to be faster, even though it executes more instructions in each iteration. However, profiling showed that this is more than compensated by fewer cache misses, probably owed to the factor two lower memory required.

### Memory reduction for backtracing and cell-off matrices

To compute an alignment by backtracing from the cell (*i*_best_, *j*_best_) with maximum score, we need to store for each cell (*i, j*) and every pair state (*MM, GD, MI, DG, IM*) the previous cell and pair state the alignment would pass through, that is, which cell contributed the maximum score in (*i, j*). For that purpose it obviously suffices to only store the previous pair state.

HHblits 2.0.16 uses five different matrices of type char, one for each pair state, and one char matrix to hold the cell-off values (in total 6 bytes). The longest known protein Titin has about 33000 amino acids. To keep a 33000 × 33000 × 6 byte matrix in memory, we would need 6 GB of memory. Since only a fraction of ~10^−5^ sequences are sequences longer than 15000 residues in the UniProt database, we restrict the default maximum sequence length to 15000. This limit can be increased with the option `-maxres`.

But we would still need about 1.35 GB to hold the backtrace and cell-off matrices. A naive SSSE3 implementation would therefore need 5.4 GB, and 10.8 GB with AVX2. Because every thread needs its own backtracing and cell-off matrices, this can be a severe restriction.

We reduce the memory requirements by storing all back-tracing information and the cell-off flag in a single byte per cell (*i, j*). The preceding state for the IM, MI, GD, DG states can be held as single bit, with a 1 signifying that the preceding pair state was the same as the current one and 0 signifying it was MM. The preceding state for MM can be any of STOP, MM, IM, MI, GD, and DG. STOP represents the start of the alignment, which corresponds to the 0 in eq. (1) contributing the largest of the 6 scores. We need three bits to store these six possible predecessor pair states. The backtracing information can, thus, be held in ‘4 + 3’ bits, which leaves one bit for the cell-off flag (Fig. 5). Due to the reduction to one byte per cell we need only 0.9 GB (with SSSE3) or 1.8 GB (with AVX2) per thread to hold the backtracing and cell-off information.

**Figure 5.**
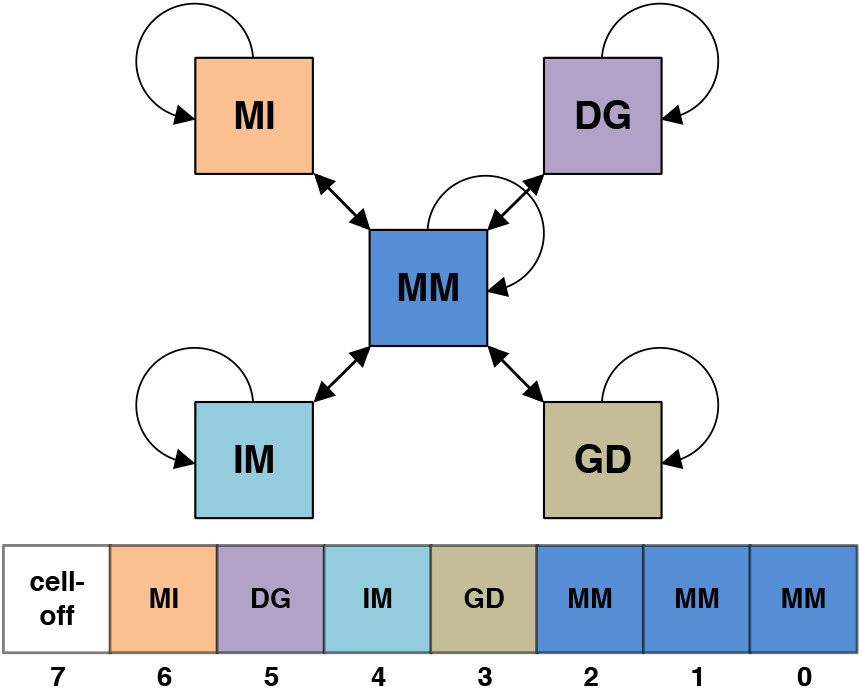
Predecessor pair states for backtracing the Viterbi alignments are stored in a single byte of the backtrace matrix in HH-suite3 to reduce memory requirements. The bits 0 to 2 (blue) are used to store the predecessor state to the MM state, bits 3 to 6 store the predecessor of GD, IM, DG and Ml pair states. The last bit denotes cells that are not allowed to be part of the suboptimal alignment because they are near to a cell that was part of a better-scoring alignment.

### Viterbi early termination criterion

For some query HMMs, the prefilter lets a lot of non-homologous target HMMs pass, for example when they contain one of the very frequent coiled coil regions. To avoid having to align thousands of non-homologous target HMMs with the costly Viterbi algorithm, we introduced an early termination criterion in HHblits 2.0.16. We averaged 1/(1 + E-value) over the last 200 processed Viterbi alignments and skipped all further database HMMs when this average dropped below 0.01, indicating that the last 200 target HMMs produced very few Viterbi E-values below 1.

This criterion requires the targets to be processed by decreasing prefilter score, while our vectorized version of the Viterbi algorithm requires the database profile HMMs to be ordered by decreasing length. We solved this dilemma by sorting the list of target HMMs by decreasing prefilter score, splitting it into equal chunks (default size 2000) with decreasing scores, and sorting target HMMs within each chunk by their lengths. After each chunk has been processed by the Viterbi algorithm, we compute the average of 1/(1 + E-value) for the chunk and terminate early when this number drops below 0.01.

### SIMD-based MSA redundancy filter

To build a profile HMM from an MSA, HH-suite reduces the redundancy by filtering out sequences that have more than a fraction `seqid_max` of identical residues with another sequence in the MSA. The scalar version of the function (Algorithm 5) returns 1 if two sequences *x* and *y* have a sequence identity above `seqid_min` and 0 otherwise. The SIMD version (Algorithm 6) has no branches and processes the amino acids in chunks of 16 (SSSE3) or 32 (AVX2). It is about ~11 times faster than the scalar version.

#### Algorithm 5 Check if x,y have seq. identity > seqid min

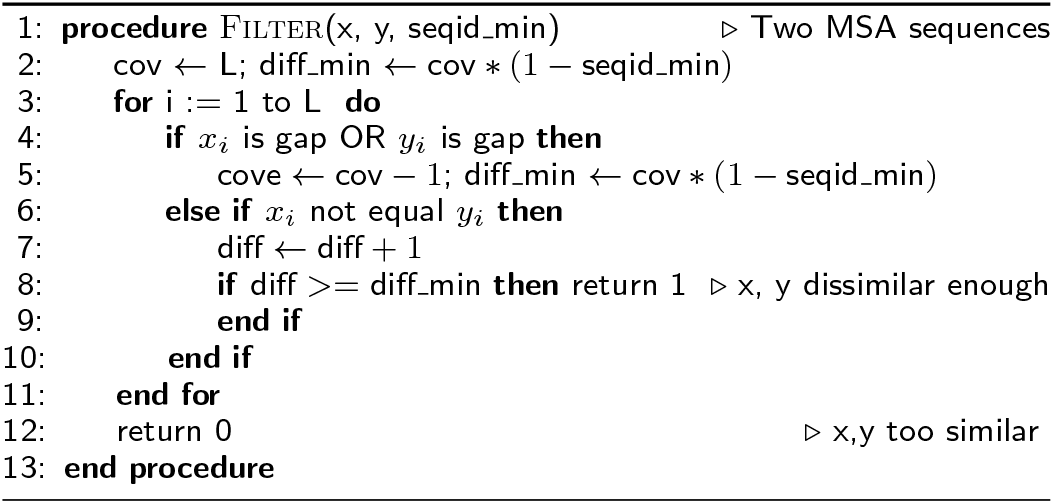

## Results

### Speed benchmarks

#### Speed of HHsearch 2.0.16 versus HHsearch 3

Typically more than 90% of the run time of HHsearch is spent in the Viterbi algorithm, while only a fraction of the time is spent in the the maximum accuracy alignment. Only a small number of alignments reach an E-value low enough in the Viterbi algorithm to be processed further. HHsearch therefore profits considerably from the SIMD vectorization of the Viterbi algorithm.

**Figure 6.**
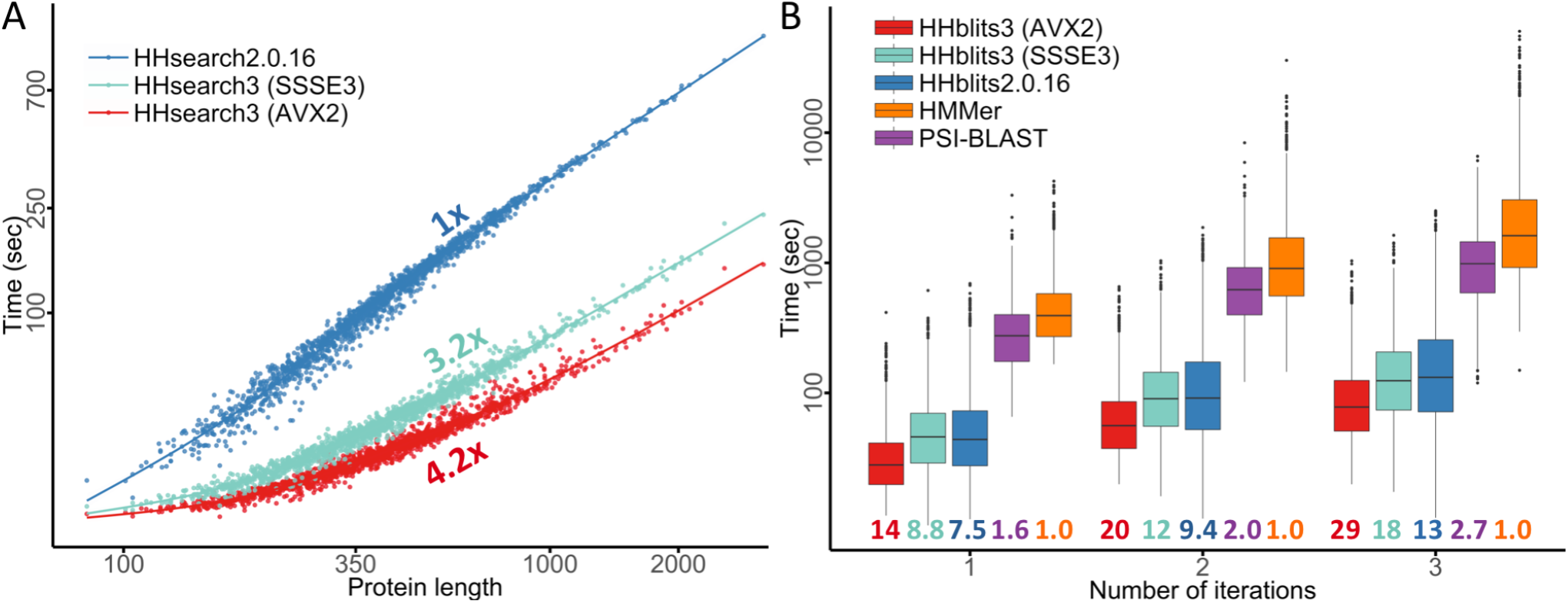
Speed comparisons. A runtime versus query profile length for 1644 searches with profile HMMs randomly sampled from UniProt. These queries were searched against the PDB70 database containing 35000 profile HMMs of average length 234. The average speedup over HHsearch 2.0.16 is 3.2-fold for SSSE3-vectorized HHsearch and 4.2-fold for AVX2-vectorized HHsearch. B Box plot for the distribution of total runtimes (in logarithmic scale) for one, two, or three search iterations using the 1644 profile HMMs as queries. PSI-BLAST and HHMER3 searches were done against the UniProt database (version 2015_06) containing 49293307 sequences. HHblits searches against the uniprot20 database, a clustered version of UniProt containing profile HMMs for each of its 7313957 sequence clusters. Colored numbers: speed-up factors relative to HMMER3.

##### Algorithm 6 Vectorized version of Algorithm 5

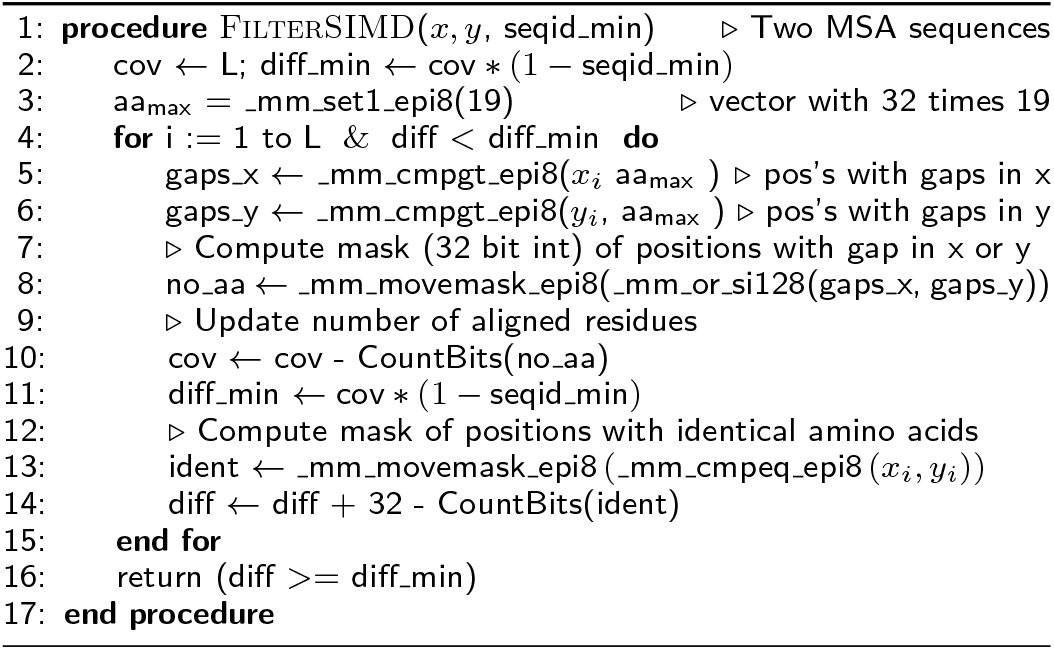

To compare the speed of the HHsearch versions, we randomly selected 1644 sequences from Uniprot (release 2015_06), built profile HMMs, and measured the total run time for searching with the 1644 query HMMs through the PDB70 database (version 05Sep15). The PDB70 contains profile HMMs for a representative set of sequences from the PDB [22], filtered with a maximum pairwise sequence identity of 70%. It contained 35 000 profile HMMs with an average length of 234 match states.

HHsearch with SSSE3 is 3.2 times faster and HHsearch with AVX2 vectorization is 4.2 times faster than HHsearch 2.0.16, averaged over all 1644 searches (Fig. 7A). For proteins longer than 1000, the speed-up factors are 5.0 and 7.4, respectively. Due to a runtime overhead of ~20s that is independent of the query HMM length (e.g. for reading in the profile HMMs), the speed-up shrinks for shorter queries.

**Figure 7.**
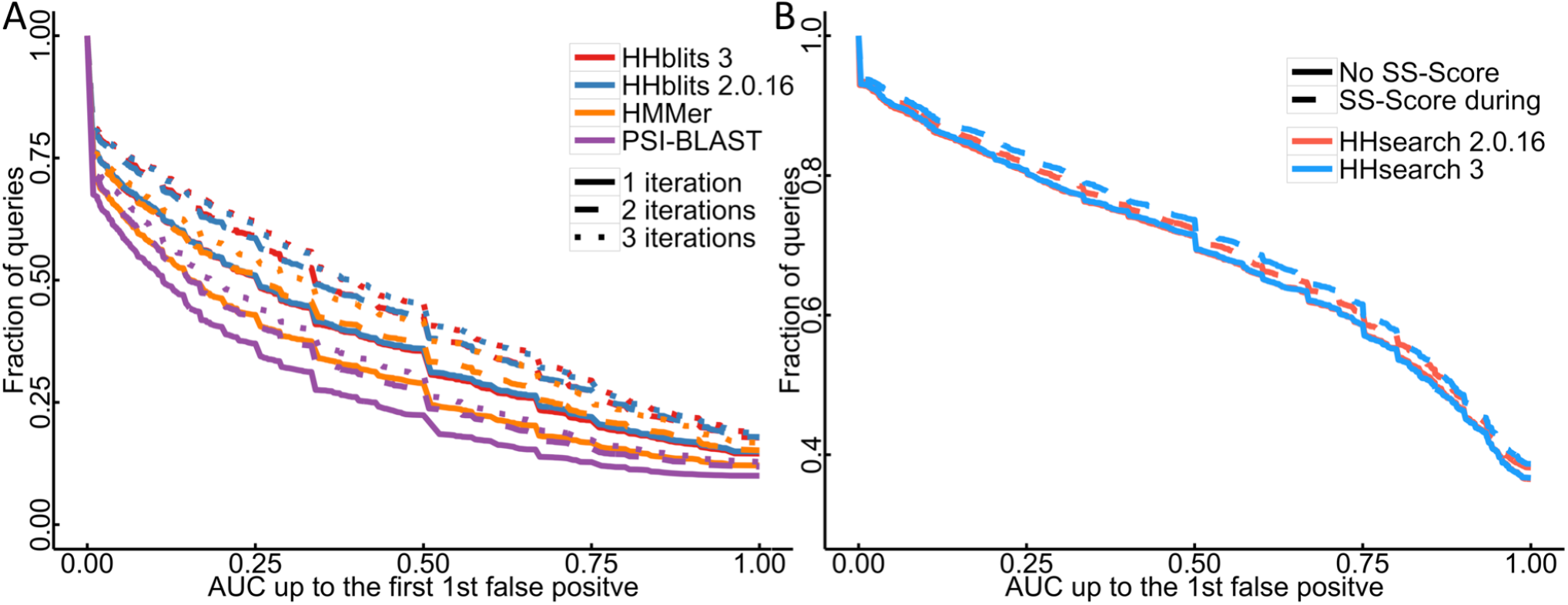
Sensitivity of sequence search tools. A We searched with 6616 SCOP20 domain sequences through the UniProt plus SCOP20 database using one to three search iterations. The sensitivity to detect homologous sequences is measured by cumulative distribution of the Area Under the Curve 1 (AUC1), the fraction of true positives ranked better than the first false positive match. True positive matches are defined as being from the same SCOP superfamily [24], false positives have different SCOP folds, excepting known cases of inter-fold homologies. B Sensitivity of HHsearch with and without scoring secondary structure similarity, measured by the cumulative distribution of AUC1 for a comparison of 6616 profile HMMs built from SCOP20 domain sequences. Query HMMs include predicted secondary structure, target HMMs include actual secondary structure annotated by DSSP. True and false positives are defined as in A.

Most of this speed-up is owed to the vectorization of the Viterbi algorithm: The SSSE3-vectorized Viterbi code ran 4.2 times faster than the scalar version.

In HHblits, only part of the runtime is spent in the Viterbi algorithm, while the larger fraction is used by the prefilter, which was already SSSE3-vectorized in HHblits 2.0.16. Hence we expected only a modest speed-up between HHblits 2.0.16 and SSSE3-vectorized HHblits 3. Indeed, we observed an average speed-up of 1.2, 1.3, and 1.4 for 1, 2 and 3 search iterations, respectively (Fig. 7A), whereas AVX2-vectorized version is 1.9, 2.1, and 2.3 times faster than HH-blits 2.0.16, respectively. AVX2-vectorized HHblits is 14, 20, and 29 times faster than HMMER3 [3] (version 3.1b2) and 9, 10, and 11 times faster than PSI-BLAST [9] (blastpgp 2.2.31) for 1, 2, and 3 search iterations.

All runtime measurements were performed using the Unix tool `time` on a single core of a computer with two Intel Xeon E5-2640 CPUs with 128 GB RAM.

### Sensitivity benchmark

To measure the sensitivity of search tools to detect remotely homologous protein sequences, we used a benchmarking procedure very similar to the one described in [4]. To annotate the uniprot20 (version 2015_06) with SCOP domains, we first generated a SCOP20 sequence set by redundancy-filtering the sequences in SCOP 1.75 [24] to 20% maximum pairwise sequence identity using `pdbfilter.pl` with minimum coverage of 90% from HH-suite, resulting in 6616 SCOP domain sequences. We annotated a subset of uniprot20 sequences by the presence of SCOP domains by searching with each sequence in the SCOP20 set with `blastpgp` through the consensus sequences of the uniprot20 database and annotated the best matching sequence that covered ≥ 90% of the SCOP sequence and that had a minimum sequence identity of at least 30%.

We searched with PSI-BLAST (2.2.31) and HMMER3 (v3.1b2) with three iterations, using the 6616 sequences in the SCOP20 set as queries, against a database made up of the UniProt plus the SCOP20 sequence set. We searched with HHblits versions 2.0.16 and 3 with three iterations through a database consisting of the uniprot20 HMMs plus the 6616 UniProt profile HMMs annotated by SCOP domains.

We defined a sequence match as true positive if query and matched sequence were from the same SCOP superfamily and as false positive if they were from different SCOP folds and ignore all others. We excluded the self-matches as well as matches between Rossman-like folds (c.2-c.5, c.27 and 28, c.30 and 31) and between the four-to eight-bladed β-propellers (b.66-b.70), because they are probably true homologs [1]. HMMER3 reported more than one false positive hit just in one out of three queries, despite setting the maximum E-value to 100000, and we therefore measured the sensitivity up to the first false positive (AUC1) instead of the AUC5 we had used in earlier publications.

We ran HHblits using `hhblits -min_prefilter_hits 100 -n 1 -cpu $NCORES -ssm 0 -v 0 -wg` and wrote check-point files after each iteration to restart the next iteration. We ran HMMER3 (v3.1b2) using `hmmsearch −chkhmm -E 100000` and PSI-BLAST (2.2.31) using `-evalue 10000 -num_descriptions 250000`.

The cumulative distribution over the 6616 queries of the sensitivity at the first false positive (AUC1) in Fig. 7A shows that HHblits 3 is as sensitive as HHblits 2.0.16 for 1, 2, and 3 search iterations. Consistent with earlier results [4, 25], HHblits is considerably more sensitive than HMMER3 and PSI-BLAST.

We also compared the sensitivity of HHsearch 3 with and without scoring secondary structure similarity, because we slightly changed the weighting of the secondary structure score (Methods). We generated a profile HMM for each SCOP20 sequence using three search iterations with HH-blits searches against the uniprot20 database of HMMs. We created the query set of profile HMMs by adding PSIPRED-based secondary structure predictions using the HH-suite script `addss.pl`, and we added structurally defined secondary structure states from DSSP [35] using `addss.pl` to the target profile HMMs. We then searched with all 6616 query HMMs through the database of 6616 target HMMs. True positive and false positive matches were defined as before.

Figure 7B shows that HHsearch 2.0.16 and 3 have the same sensitivity when secondary structure scoring is turned off. When turned on, HHsearch 3 has a slightly higher sensitivity due to the better weighting.

## Conclusions

A sizeable fraction of proteins in genomics and metagenomics projects remain unannotated due to the lack of an identifiable, annotated homologous protein [36]. These - projects could improve their annotation depth by adding HHblits searches through the PDB, Pfam and other alignment databases to their annotation pipeline [7] at only a marginal cost in CPU time, since HHblits 3 runs 20 times faster than HMMER, the standard tool for Pfam and INTERPRO annotations. The improvements in parallelization across CPU cores and compute servers further facilitates the use of HH-suite on larger projects.

## Availability and requirements

- Project name: HH-suite
- Project page:https://github.com/soedinglab/hh-suite
- Operating systems: Linux, Max OS X
- Programming languages: C++ with SSSE3/AVX2 intrinsics, Python utilities
- Other requirements: support for SSSE3 or higher
- License: GNU GPL version 3

## Abbreviations

MSA: :multiple sequence alignment;
HMM: :hidden Markov model;
SIMD: :single-instruction multiple-data;
SSSE3: :supplemental streaming SIMD extensions 3;
AVX2: :advanced vector extension (SIMD instruction set standards);

## Declarations

### Ethics approval and consent to participate

Not applicable

### Consent for publication

Not applicable.

### Availability of data and materials

The datasets used and/or analysed during the current study are available from the corresponding author on request.

### Author contributions

MS & JS designed research, MS developed vectorized code and performed analyses, M. Meier refactored code, added features, fixed bugs and performed benchmarks, M. Mirdita added features, fixed bugs and maintains databases, HV implemented mmCIF support, SH optimized the MAC algorithm memory usage, MS and JS wrote the manuscript.

